# Temperature and intraspecific variation affect host-parasite interactions

**DOI:** 10.1101/2023.08.24.554680

**Authors:** Sherine Ismail, Johannah Farner, Lisa Couper, Erin Mordecai, Kelsey Lyberger

## Abstract

Parasites play key roles in regulating aquatic ecosystems, yet the impact of climate warming on their ecology and disease transmission remains poorly understood. Isolating the effect of warming is challenging as transmission involves multiple interacting species and potential intraspecific variation in temperature responses of one or more of these species. Here, we leverage a wide-ranging mosquito species and its facultative parasite as a model system to investigate the impact of temperature on host-parasite interactions and disease transmission. We conducted a common garden experiment measuring parasite growth and infection rates at seven temperatures using 12 field-collected parasite populations and a single mosquito population. We find that both free-living growth rates and infection rates varied with temperature, which were highest at 18-24.5°C and 13°C, respectively. Further, we find intraspecific variation in peak performance temperature reflecting patterns of local thermal adaptation—parasite populations from warmer source environments typically had higher thermal optima for free-living growth rates. For infection rates, we found a significant interaction between parasite population and nonlinear effects of temperature. These findings underscore the need to consider both host and parasite thermal responses, as well as intraspecific variation in thermal responses, when predicting the impacts of climate change on disease in aquatic ecosystems.

## Introduction

Climate change is predicted to have widespread impacts on the dynamics and distributions of diseases, including those caused by parasites responsible for human illness (e.g., malaria), wildlife declines (e.g., chytridiomycosis in frogs), and agricultural losses (e.g., wheat stripe rust) (Smith et al. 2014). Disease outbreaks in many systems have increased as temperature conditions have become warmer overall, more extreme, and more variable (Harvell et al., 2009; Cohen et al., 2020). In recent decades, there have been sustained calls to incorporate thermal biology into disease ecology (Blanford and Thomas, 1999; Harvell et al., 2002; Lafferty, 2009; Rohr et al., 2011; Claar and Wood, 2020). A key principle established by the large body of research in response to these calls is that the hump-shaped relationship to temperature seen in many physiological processes extends to disease thermal responses (Molnár et al., 2017; Mordecai et al., 2019). Specifically, infections increase with temperature up to a peak, and then decline rapidly as temperatures exceed the thermal optimum. These thermal performance curves (TPCs) can be used to make predictions about when and where temperature conditions are favorable for disease transmission (Ryan et al., 2015, 2019; Rose et al., 2016). Because TPCs for most host - parasite systems are measured only for single populations, the extent to which parasitism thermal responses vary among populations is poorly understood, and studies assessing climate suitability for disease transmission assume consistent thermal responses (Sternberg and Thomas, 2014). This knowledge gap is of critical relevance to understanding climate change impacts on host – parasite interactions both at large spatial scales and for specific populations of conservation concern.

Many studies have investigated how temperature affects single populations of hosts, parasites, and their interactions with one another. Prior work has shown that temperature impacts parasitism through its combined effects on key host and parasite traits that determine the magnitude and timing of contact rates and interaction outcomes, such as host and parasite growth and development, host immune defenses, and parasite replication (Gehman et al., 2018; Shocket et al., 2018; Lafferty and Mordecai 2016; Mordecai et al., 2019). For example, experiments in aquatic and marine trematode parasite systems showed that high temperatures decreased parasite transmission by affecting host and parasite development rates in ways that reduced temporal overlap between infected primary intermediate hosts and susceptible definitive or secondary intermediate hosts (Paull and Johnson, 2014; Díaz-Morales et al., 2022). Current scientific understanding of host - parasite thermal ecology also recognizes the importance of sources of natural complexity such as temperature fluctuations and host microbiomes, both of which have been shown to alter host - parasite interaction outcomes (Palmer-Young et al., 2019; Kunze et al., 2022).

Despite evidence that intraspecific variation can be substantial and have significant ecological consequences, the extent of thermal response variation within host – parasite systems remains a knowledge gap, particularly for parasites and across large spatial scales (Bolnick et al. 2011, Hector et al. 2021, Wood et al. 2022). Most prior studies addressing thermal response variation in host - parasite systems examined variation within host populations when exposed to a single parasite source. Sympatric Daphnia host lineages, diatom host genotypes, and salamander host color morphs all differed in their susceptibility to bacterial or fungal parasites across temperatures (Mitchell et al. 2005, Gsell et al. 2013; Venesky et al. 2022). One study of differences among parasite populations found that two nematode forest pathogen populations from different climates had similar thermal niches when grown in culture (Pimentel and Ayres 2018), but TPCs for parasites in isolation do not consistently predict the temperature dependence of host – parasite interactions (Cohen et al., 2017). Further, significant intraspecific variation in TPC’s has been found along geographic gradients (reviewed in Tuzun and Stoks 2018). Therefore, quantifying the temperature dependence of parasitism for parasite populations from across species’ ranges can provide new insight into climate impacts on disease.

In this study, we use a larval mosquito and its aquatic parasite to understand range-wide variation in parasite thermal performance. The western tree-hole mosquito, *Aedes sierrensis*, and its facultative ciliate parasite, *Lambornella clarki*, inhabit water-filled tree cavities in western North America, which serve as natural microcosms for host-parasite interactions. This system is ideal for addressing questions about the ecology of host - parasite interactions in part because, like other aquatic ecosystems, water-filled treeholes are self-contained and replicated across landscapes, the host and parasite are geographically widespread, and infections are frequently fatal (Washburn et al. 1986, 1988). Additionally, while empirical research on thermal performance in host - parasite interactions is often limited by the difficulty of rearing participant species in isolation under lab conditions (Molnár et al. 2017), both Ae. sierrensis and L. clarki can be reared together or separately under laboratory conditions. *L. clarki* belongs to a group of free-living ciliate parasites detected in 20 mosquito species (Corliss and Coats 1976). The range of *Ae. sierrensis* and *L. clarki* spans a large geographic and temperature gradient, ranging from southern California to northern Oregon (Hawley 1985, Washburn and Anderson 1986, Broberg and Bradshaw 1997). A previous field survey found that parasite prevalence generally increased with latitude, suggesting that cooler temperatures may promote higher infection rates due to effects on the parasite or host-parasite interactions (Washburn and Anderson 1986). Importantly, because *L. clarki* is a facultative parasite that can grow and complete its life cycle in a free-living (“trophont”) stage, we can independently assess the temperature-dependence of free-living parasite growth versus parasite infectivity. Additionally, host - parasite interactions can be easily tracked because both infection and the host immune response of melanization are visible in live larvae.

We aim to quantify temperature-dependent infection rates and parasite thermal adaptation using 12 populations of parasites and a single mosquito population, replicated across seven temperatures. Specifically, we test the following three hypotheses. First, we hypothesize that because ectothermic organismal responses vary nonlinearly with temperature, infection rates will peak at intermediate temperatures that optimize parasite growth versus host defense. Second, we hypothesize that infection rates will vary across parasite populations because they span a large geographic and climatic gradient, and similar intraspecific variation is well-documented in other systems. Third, we hypothesize that both free-living and parasitic ciliates will show a signature of local adaptation to temperature (*i.e.*, parasites from colder environments will have a lower optimal temperature for growth and infection than those from warmer environments).

## Methods

### Host and parasite life cycles

*Aedes sierrensis* spans western North America from southern California to Canada and from coastal to montane environments. *Lambornella clarki* has been documented across a similar gradient from southern California to Oregon. These ranges span a substantial gradient in temperature and precipitation patterns. Following significant rainfall in fall, winter, and/or spring (depending on the location), tree holes fill with water and mosquito larvae and *L. clarki* hatch from their desiccation-resistant dormant states. At higher latitudes and elevation, larvae may go into diapause during colder months (Jordan and Bradshaw 1978). Mosquitoes transition through four larval stages, a pupal stage, then emerge as adults before the treeholes dry out during the spring and summer. A previous study conducted over a similar geographic area demonstrated temperature-dependent life history variation among *Ae. sierrensis* populations (Couper et al. 2023). Infection typically occurs during larval stages. The mosquito larvae feed on microorganisms, including free-living *L. clarki*, and detritus. In tree holes with larvae, *L. clarki* can transform into parasitic forms within 24 to 48 hours, whereas in tree holes without larvae they live as free-living ciliates, feeding on microbes in the water (Washburn et al. 1988).

### Mosquito Source

We chose one population of *Ae. sierrensis* mosquitoes to infect with many populations of the *L. clarki* parasite. To produce the larvae for this experiment, we propagated (as follows) the first generation of offspring from larvae collected from Jasper Ridge Biological Preserve (37.4086923, −122.233876) in San Mateo County, California, USA, on multiple occasions from October to December 2021. Larvae were maintained in plastic containers and fed with a 4% solution of 50% high protein cat chow, 36% bovine liver powder, and 14% brewers yeast (Maïga 2017). Upon pupation, adults were transferred to aluminum collapsible cages (BioQuip, Rancho Dominguez, CA, USA). Once emerged, adults were fed with a 10% sugar-water solution twice a week and blood-fed once a week with defibrinated sheep’s blood delivered through a membrane feeding system (Hemotek, Blackburn, UK). Oviposition cups, which consisted of a paper cup lined with damp filter paper, were placed inside each cage. Cages were checked at least once a week for eggs, and if found, they were collected and maintained at 21°C for 14 days. These eggs were then stored in the dark for 24 hours at 5°C for a minimum of 14 days to elicit diapause before being used in the experiment. Rearing methods were based on Solano County Mosquito Abatement District protocols (pers. comm., B. Barner).

### Parasite Sources

We collected larval *Ae. sierrensis* mosquitoes from field sites across California and Oregon to obtain individuals infected with *L. clarki*. This survey covered the entire known range of *L. clarki,* spanning 1500 km (Washburn and Anderson 1986, Broberg and Bradshaw 1997). Field-collected larval mosquitoes were maintained at 7°C and 24-hour dark conditions during transport and in the lab until visual inspection for *L. clarki* infections. We successfully cultured *L. clarki* from infected mosquitoes found at 11 distinct geographic sites, and at one site —in Marin County, California—we cultured two populations of *L. clarki* from two different tree holes. These infected populations, including the two tree holes from Marin, were used in our experiment for a total of 12 *L. clarki* populations (Table S1, Figure 1). In order to isolate the parasite, infected mosquitoes were crushed and placed in vials with 1 mL of autoclaved culturing media (made up of 1 L distilled water, 1 protozoan pellet, and 0.38g of Herptivite) and an autoclaved barley seed (Fukami, 2004). These cultures were maintained at 21°C and fresh media was added weekly until the experiment began.

**Figure 1.**
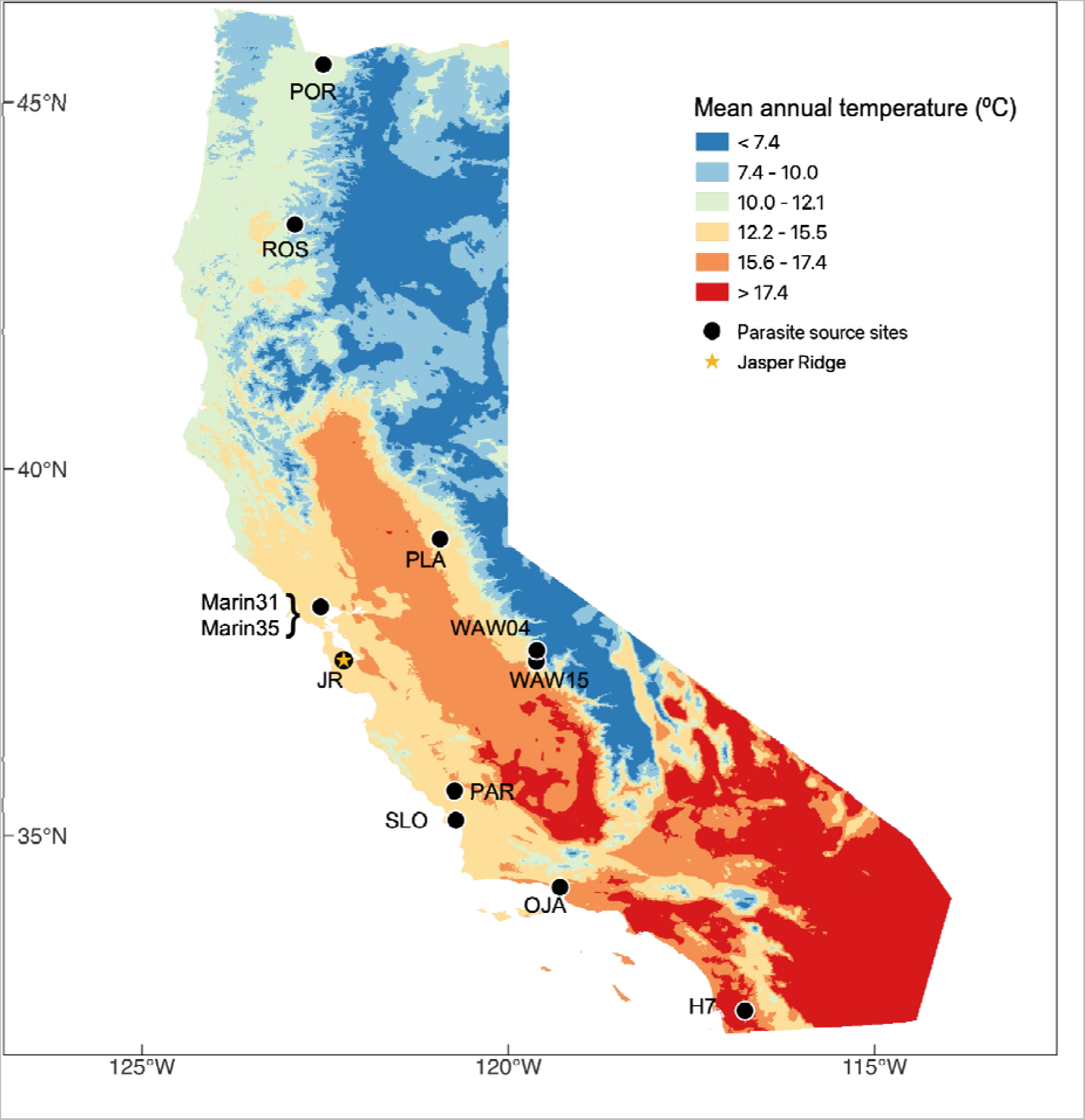
Parasites were collected across a large geographic and temperature gradient. A map of the twelve *L. clarki* parasite populations in California and western Oregon, where colors represent the mean annual temperature. The star symbol indicates the location of the mosquito source, Jasper Ridge Biological Preserve.

**Figure 2.**
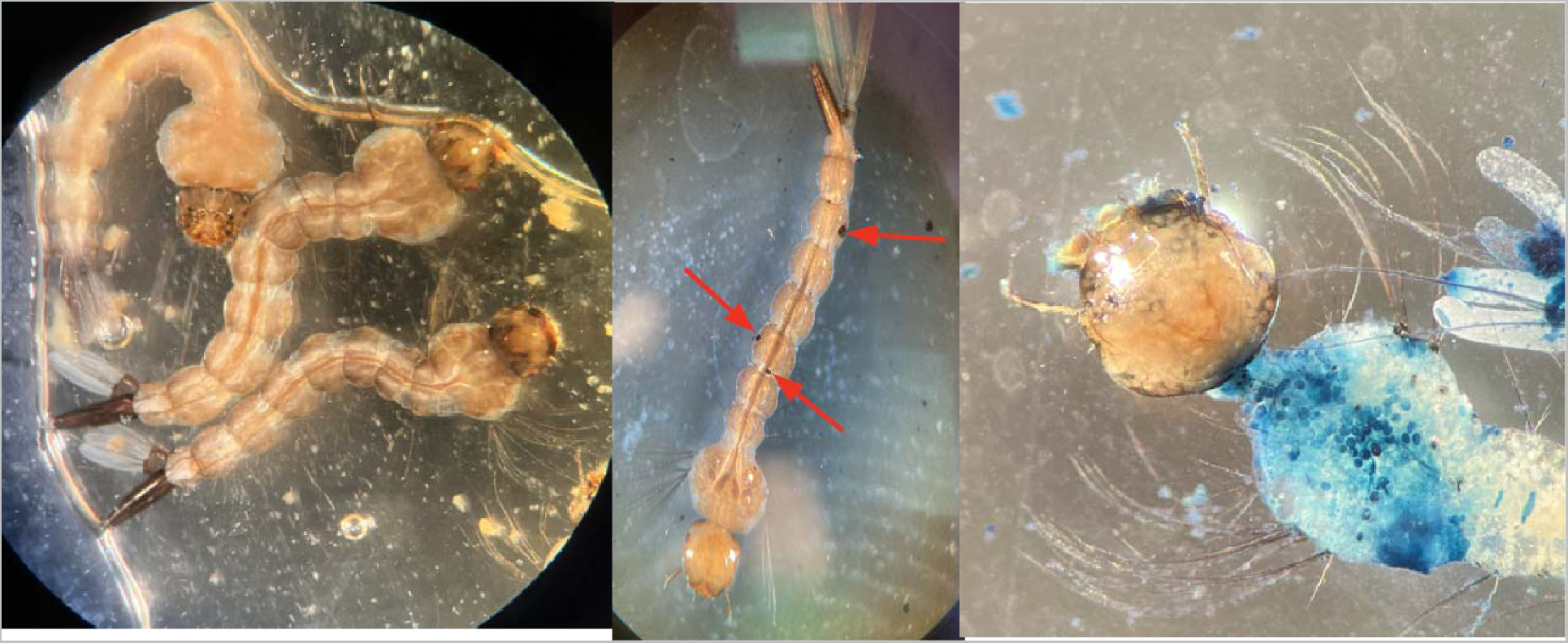
Mosquito larvae exhibit signs of infection with *L. clarki*. Experimental larval mosquitoes that are uninfected (left), exhibit melanization spots (center: red arrows), and infected with *L. clarki* cysts and cells, stained with amide dye (right: blue and white spots, respectively).

To obtain a measure of natal source temperature from our parasite collection sites, we extracted the 19 bioclimatic variables from the WorldClim dataset (Fick and Hijmans 2017). Each variable is available at a 1 km^2^ resolution, and we use the annual averages from 1997 - 2000. We focus on the mean annual temperature when assessing thermal adaptation (Table S1). Although hosts and parasites are not active after tree holes have dried out in late summer, this ensures that we capture their active period across the large geographic extent of our study. The mean annual temperature ranged from 11-18°C at our collection sites.

### Infection Experiment

We examined geographic patterns in the temperature-dependence of parasite infectivity across a thermal gradient through a common garden experiment performed in the laboratory. In this experiment, we used larvae hatched from a single location (*i.e*., Jasper Ridge) (Figure 1). Host eggs were hatched synchronously by immersion in a solution made up of 500mL Arrowhead distilled water, 300mL autoclaved tree hole water, and 1 teaspoon Brewer’s yeast (Schwan & Anderson, 1980).

Freshly hatched, first-instar larvae (less than 24 hours old), were isolated into groups of five. We pipetted 5 larvae in 1 mL of the hatching solution into each well of a 6-well non-treated Falcon plate. For each *L. clarki* population, we counted free-living cells using a dissecting microscope and diluted the cultures with fresh media to establish a standard *L. clarki* inoculation concentration of 500 cells/mL. This dose was chosen based on pilot experiments in which much lower densities led to no infection and much higher densities led to infection in nearly all mosquitoes. We added 4 mL of this concentration to each well of five larvae. Once all wells were prepared, we placed the trays at one of seven constant temperatures: 5, 9, 13, 17, 21, 24, and 28°C. To maintain temperature treatments, trays were placed in styrofoam boxes housed inside 7 incubators, one for each temperature. In total, we had three replicates for each *L. clarki* population by temperature treatment. These replicates were maintained at 24h darkness and fed three times per week with 25 uL of 4% larval food suspension, equating to 0.11 mg food per larva per day, which was chosen so that resources would not limit mosquito growth (Maïga 2017). The experiment ended when the first larva of a replicate began to pupate, which lasted a total of 30 days at the coldest temperature, and fewer days at warmer temperatures; thus, temperature inherently affects exposure time.

We examined each larva twice a week using a dissecting microscope at 40X magnification. During the examinations, we recorded infections, melanization spots—an immune response to infection—and survival. As a final assay of infection, dead larvae were stained for 30 minutes with black amide dye to ascertain whether *L. clarki* cuticular cysts were present on the host’s cuticle (Soldo & Merlin, 1972, Washburn et al., 1988). Similarly, once the first pupa was observed at a given temperature, all 4th instar larvae at that temperature were stained with amide dye as an additional check for cuticular cysts, which are a sign of cuticular infection, and were thus considered to be an infection in our experiment.

### Free-living Ciliate Thermal Performance Experiment

Since *L. clarki* is a facultative parasite, we could measure the growth rates of free-living cells. To determine the thermal performance of the parasites in the absence of the host, we measured growth rates of free-living *L. clarki* across the same suite of temperatures as in the infection experiment: 5, 9, 13, 17, 21, 24, and 28°C, maintained in the same incubators as the infection experiment. We began with a 2 mL solution with a concentration of 3 cells/100 uL with 5 replicates of each of the 12 populations. We then measured population density daily for 4 days by visually counting individuals in 100uL subsamples using a dissecting microscope at 40X magnification.

### Statistical analysis

We classified larvae that had either visible internal infections or cuticular cysts as infected and larvae that lacked these signs as uninfected. We used the number of infected larvae per replicate of 5 larvae as our dependent variable. To examine the effect of temperature, and to determine whether there was significant variation in infection among parasite populations, we ran a Poisson generalized linear regression using the “glm” function in the “stats” package in R version 4.1.1 (, R Core Team 2021). We use this model as our data was approximately Poisson distributed and Poisson models are capable of effectively handling low infection counts. Because we expected infection to have a nonlinear response to temperature that might vary across populations, our full model included temperature, temperature squared, *L. clarki* population, the interaction between *L. clarki* population and temperature, and the interaction between *L. clarki* population and temperature squared. We did not include tray as a fixed effect as the full model including tray failed to converge. We checked model assumptions using the “simulateResiduals” function in the “DHARMa” package (Hartig 2017). The q-q plot and plot of observed versus predicted residuals showed no signs of significant patterns or deviations. We report significance using likelihood ratio tests. We also investigated variation in infectivity based on mosquito immune responses, following the same process as above but using the number of larvae with melanization as the dependent variable. As before, we performed model selection and determined significance using likelihood ratio tests.

To analyze the free-living ciliate thermal performance data, we calculated exponential growth rates for each replicate using nonlinear least squares (“nls”) in R. All further analysis was done using the “rTPC” package in R (Padfield et al. 2021). For each population, we fit a thermal performance curve using the “sharpeschoolhigh_1981” model, which describes a left-skewed curve (Schoolfield et al. 1981). We calculated the curve breadth as the range of temperatures over which a curve’s rate is 80% of peak using “get_breadth” and the activation energy parameter as estimated by the Sharpe-Schoolfield model. We bootstrapped using “Boot” to produce 95% confidence intervals. We considered non-overlapping confidence intervals as significant differences between populations. To test for local adaptation, we conducted a weighted linear regression of peak performance temperature as a function of mean annual temperature at the source of the population, with weights equal to the inverse variance of bootstrapped estimates, which allows us to propagate uncertainty around the peaks. We also calculated Pearson’s correlation coefficients between the 19 bioclimatic variables and peak performance temperature.

## Results

### Mosquito-Parasite Infection Experiment

We observed infections at all temperatures and in all parasite populations. When averaging across populations, infection rates were highest at intermediate temperatures (13°C) and were lowest at 21°C and 24°C (Figure 3A). Some parasite populations showed a clear hump-shaped relationship between infection and temperature (*e.g.*, Marin31, Marin35, and ROS07), while others showed no clear trend (*e.g.*, WAW15, PLA, and PAR; Figure 3B). Consistent with our hypothesis, the results of our generalized linear regression analysis revealed a quadratic relationship between temperature and infection. Temperature was significant (χ^2^ = 7.70, df = 1, p = 0.01) as was temperature squared (χ^2^ = 12.17, df = 1, p < 0.001). Further, our analysis demonstrated that parasite populations exhibited distinct responses to temperature. Although there was no significant effect of population alone, there was a significant interaction between population and temperature (χ^2^ = 20.44, df = 11, p = 0.04) and between population and temperature squared (χ^2^ = 26.48, df = 11, p = 0.006). These findings reveal that the relationship between infection and temperature is influenced by the specific population being studied.

**Figure 3.**
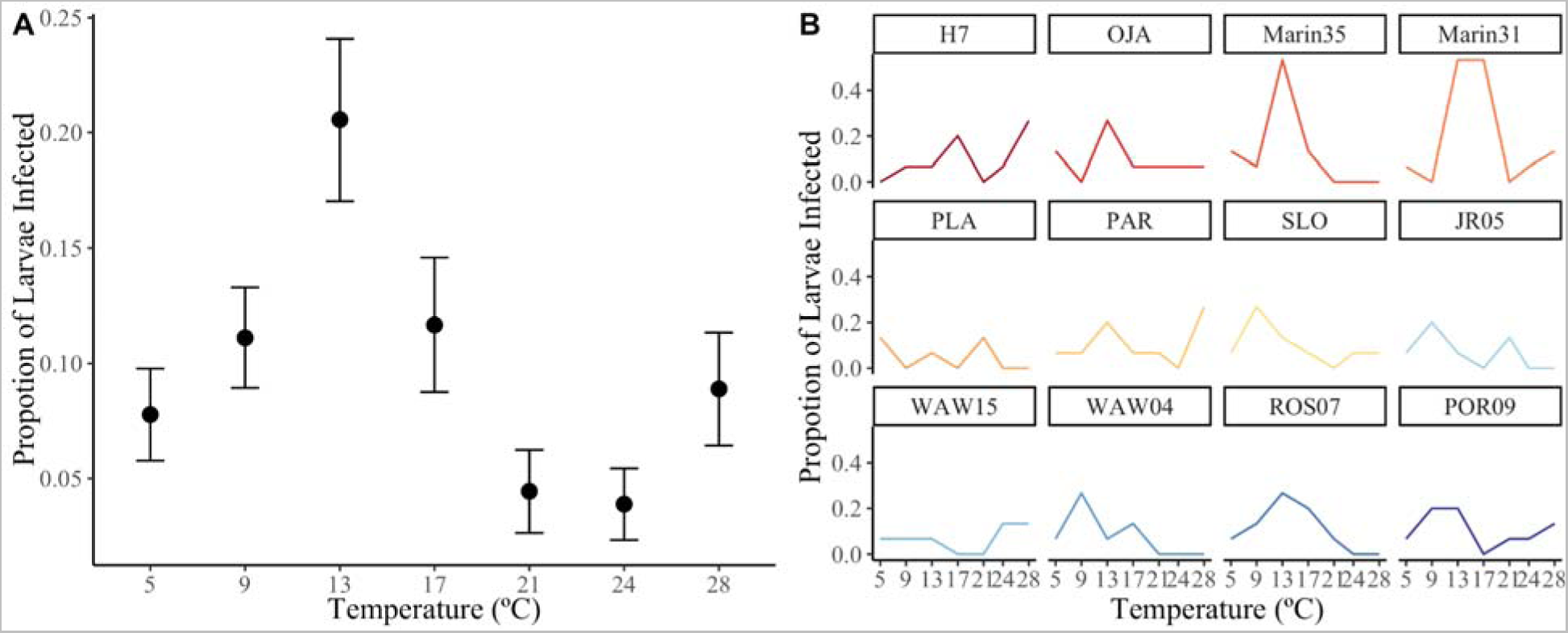
Infections peaked at 13°C, and this thermal response is population specific. A. Proportion of larvae infected (±1 SE) averaged across all parasite populations, shown for each temperature. Infections peaked at 13°C. B. Average proportion of larvae infected across temperature for each population. Populations are colored in order of decreasing source temperature, defined as mean annual temperature.

Overall, the Marin31 population had the greatest average number of observed infections (19% infected), while PLA had the lowest (5% infected) (Figure 4). SLO, OJA, and H7 populations had similar, intermediate rates of infections with an average of 10% infected. Despite both populations being from the same location, Marin31 and Marin35 had different infection rates (19% infected in Marin31 compared to 12% infected in Marin35), although overall, populations were not found to be significantly different.

**Figure 4.**
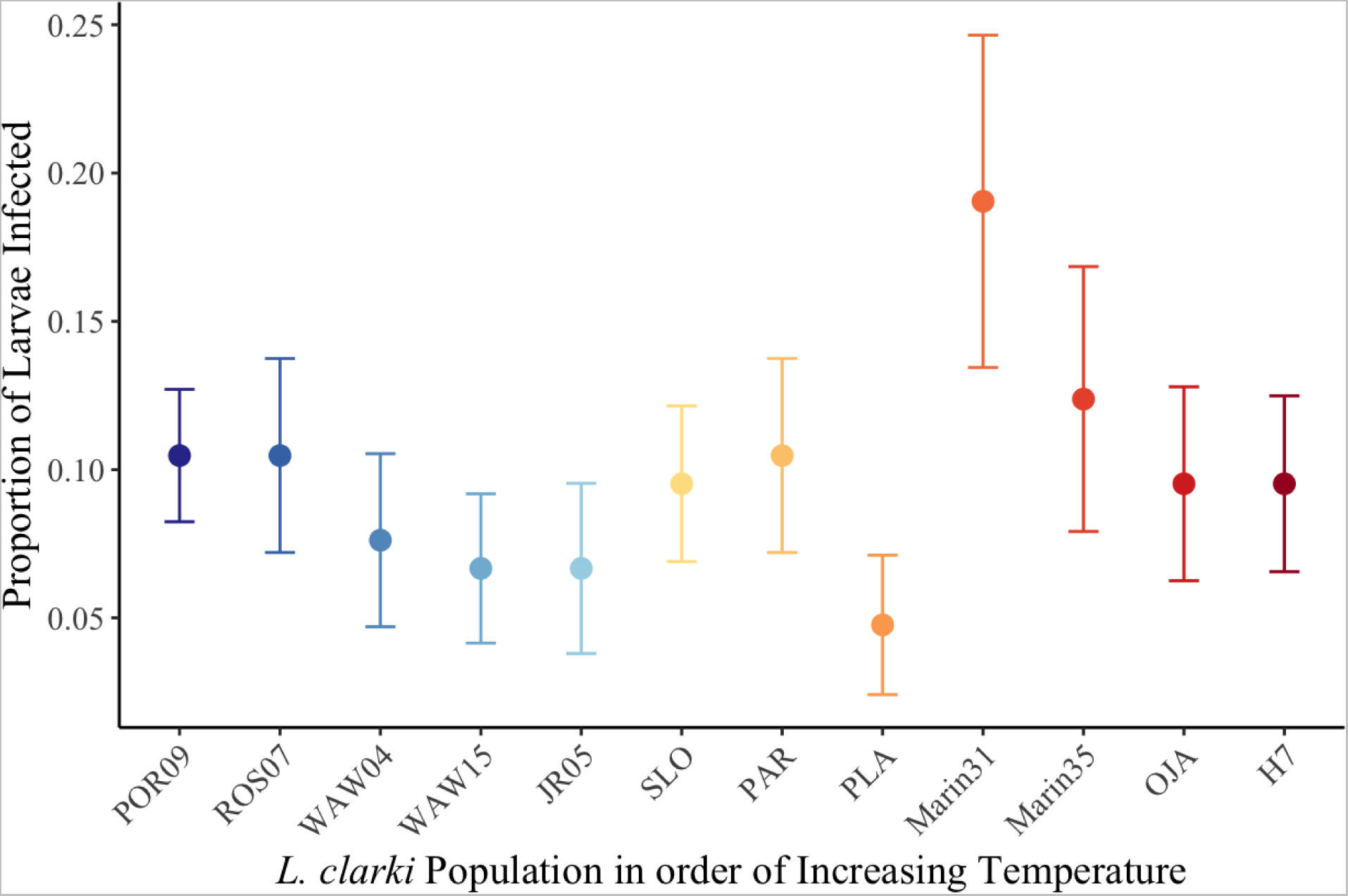
Populations varied in infection rates but not systematically with source temperature. Proportion of larvae infected (±1 SE) averaged across all temperatures, shown for each of the 12 parasite populations. Populations are in order of increasing source temperature, defined as mean annual temperature.

For melanization rates, we found that the parasite population affected the number of larvae launching an immune response (i.e., the number melanized). The model results show differences in melanization rates among parasite populations (χ^2^ = 24.55, df = 11, p = 0.01). There was also a significant interaction between population and temperature (χ^2^ = 20.21, df = 11, p = 0.04) but not between temperature and temperature squared. We observed the highest number of larvae melanized at 13°C and no melanization at or above 21°C (Figure S1).

### Free-Living Ciliate Thermal Performance Experiment

Across all populations, we observed a nonlinear, left-skewed relationship between temperature and *L. clarki* growth rate in the absence of the host (Figure 5; Figure S2). Ciliate growth rates peaked between 18.0–24.5°C (Table S1) and declined steeply above the peak. Further, we found that ciliate populations differed in their thermal performance curves. Populations differed with respect to their breadth (range = 6.38–15.84°C, mean = 9.02, sd = 2.57), and activation energy (range = 0.26–0.69 eV, mean = 0.49, sd = 0.11) in addition to peak performance temperature. For example, WAW04 and JR had wide performance breadths compared to other populations (Figure 5). Notably, Marin31 had a steep decline beyond 25°C, which was driven by all replicates in the 28°C treatment going extinct by the second day (Figure 5).

**Figure 5.**
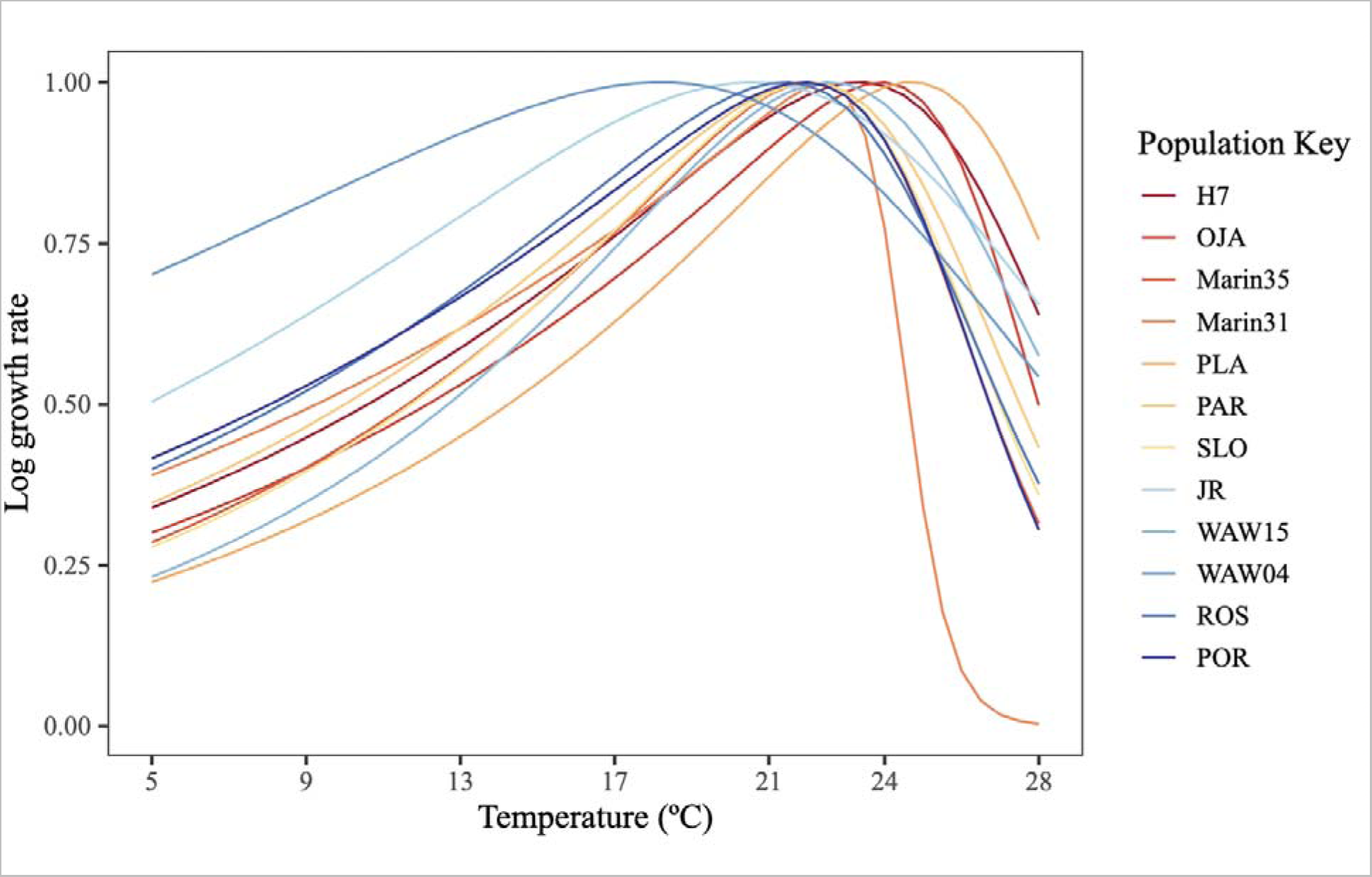
Free-living ciliate growth varies nonlinearly with temperature and across populations. Thermal performance curves of free-living parasite population growth rates fit using a Sharpe-Schoolfield model. Growth rates are measured in units of cells per 100ul per day. Populations are colored in order of decreasing source temperature, defined as mean annual temperature. WAW04 is an outlier at colder temperatures and Marin31 is an outlier at warmer temperatures.

We found that populations from colder climates had lower peak performance temperatures compared to those from warmer climates, suggesting that *L. clarki* may be thermally adapted, although this relationship was marginally significant (β = 0.40, se = 0.20, p = 0.07; Figure 6). We also found positive correlations between peak performance temperature and temperature-related bioclimatic variables, including but not limited to mean annual temperature (r = 0.65) and wettest quarter temperature (r = 0.52, Table S2). The population from the coldest source temperature, WAW04, had the lowest peak performance temperature of 18°C, although this population also had the highest uncertainty around this estimate (Figure 6). WAW04 is a high elevation Sierra Nevada site, which could expose this population to qualitatively different, colder winters than sites with similarly cold annual temperatures at high latitudes. The population from the warmest temperature and lowest latitude in Southern California, H7, had the second-highest peak performance temperature of 24°C (Figure 6). In contrast to this general trend, PLA had the highest peak performance temperature of 25°C despite being collected from a relatively cold location in the Sierra Nevada foothills (Figure 6).

**Figure 6.**
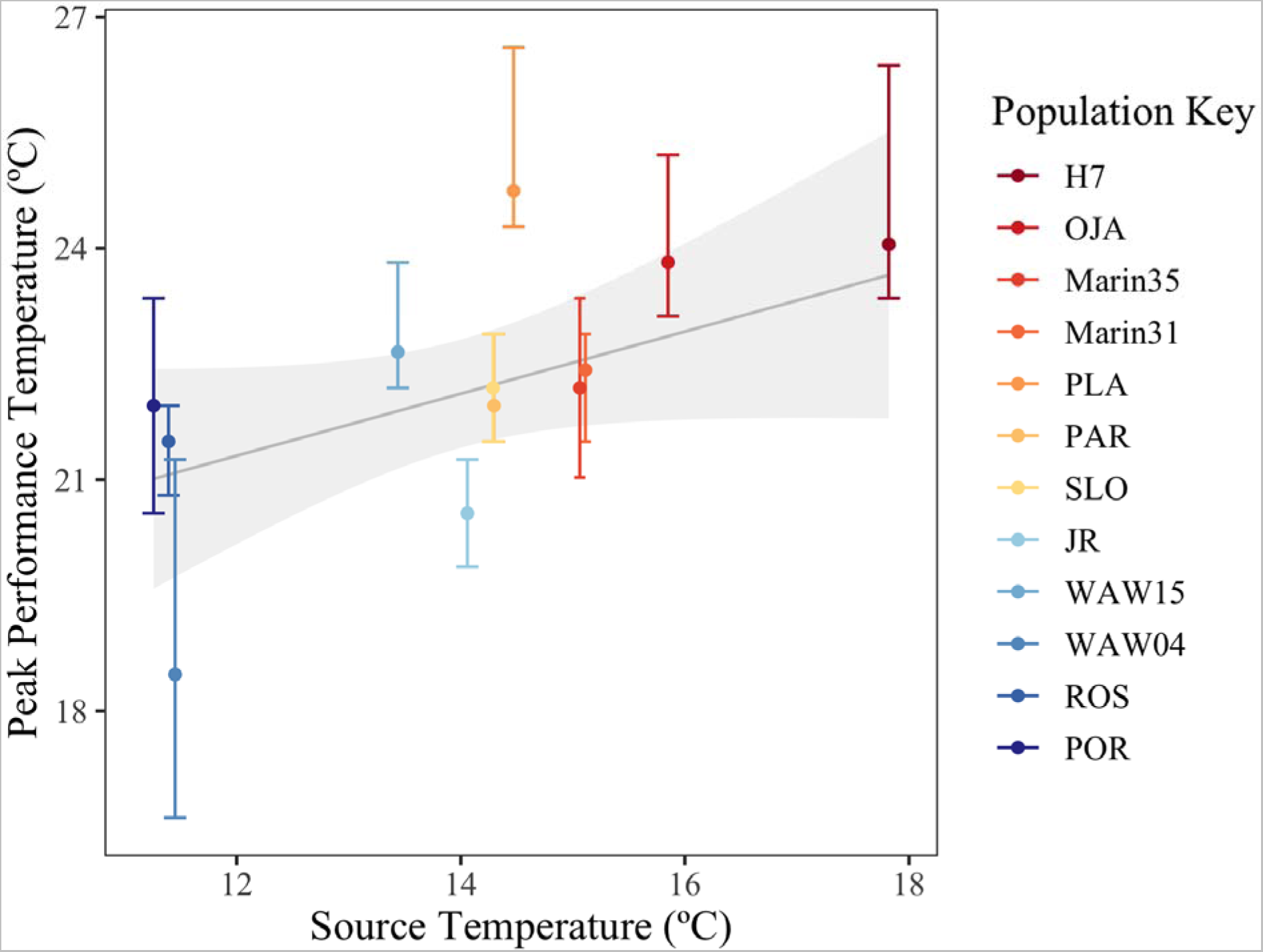
Temperature of peak free-living growth has a positive relationship with temperature in the source environment, consistent with thermal adaptation. Mean and 95% confidence interval of peak performance temperatures of ciliate populations (points and error bars). Populations are colored in order of decreasing source temperature, defined as mean annual temperature. The regression line and 95% confidence interval is in gray.

## Discussion

Temperature can have a drastic impact on host-parasite interactions (Lafferty et al. 2009, Rohr et al. 2011, Gehman et al. 2018). However, despite previous findings that intraspecific variation can be substantial (Bolnick et al. 2011, Wood et al. 2022) and can mediate range-wide responses to climate change (Gilman et al. 2006, DeMarche et al. 2019, Herrando-Pérez et al. 2019), most studies on the effects of temperature on parasitism focus on a single population of host and parasite. In this study, we examined the temperature-dependent growth and infection rates of 12 populations of a mosquito parasite from across its known geographic range (Figure 1). Our results suggest that free-living ciliate population growth is adapted to local climate: there was a marginally significant positive relationship between the thermal optima for population growth and the mean annual temperature in the source environment (Figure 6). Our infection experiment showed a significant interaction between population and thermal response, although the observed variation in thermal response did not indicate a discernable pattern of thermal adaptation. We also found that infection rates have a quadratic relationship with temperature, peaking at an intermediate temperature of 13°C (Figure 3), well below the optimal temperatures of 18.0– 24.5°C for ciliate growth rates in the absence of the host (Figure 5).

Several mechanisms may help explain the difference in peak infection temperature and peak free-living ciliate growth temperature. First, the contrasting temperatures may be due to a temperature-sensitive immune response in the host, which could reduce parasite infection at temperatures that optimize the immune response. Murdock et al. (2012a) found that melanization, a non-specific immune defense, peaked at 18°C in response to the injection of non-pathogenic particles in the Asian malaria mosquito, *Anopheles stephensi*. Additionally, in insects more generally, warmer temperatures can increase phenoloxidase activity, an important enzyme in the melanization pathway (Murdock et al. 2012b). In our experiment, melanization occurred in response to cuticular infection, so we were not able to disentangle the direct effect of temperature on melanization from its effects on infection. Second, warmer temperatures accelerate larval development rates, which limits the timeframe for possible infections to occur. We observed this phenomenon in our experiment—the first day of pupation declined with temperature from day 30 at 9°C to day 9 at 28°C, and no individuals pupated at 5°C (Figure S3). Increased larval development rates also result in faster molting—transitions between the larval instar stages— which allows larvae to shed infective cysts attached to their cuticle. This increased rate of molting may also have been responsible for our failure to detect melanization spots that formed on the cuticle at temperatures above 17°C (Figure S1), as we only checked larvae twice a week. Therefore, the differences in peak infection temperature and peak free-living growth temperature are likely due to the temperature-sensitive immune responses in the host, and accelerated larval development rates that limit the timeframe for infections.

The effects of warmer temperatures on infection ultimately depend on the relative thermal sensitivities of host and parasite traits. In this system, temperature-dependent host melanization and *L. clarki* growth rates are two such traits driving infection dynamics. One theoretical framework for incorporating these thermal sensitivities into a prediction is the thermal mismatch hypothesis. This hypothesis posits that parasitism rates often peak at temperatures that are relatively less suitable for the host than the parasite. Specifically, if thermal performance curves of both host and parasite are left-skewed, peak infection should occur at cool temperatures (and vice versa for species with right-skewed curves), because parasites typically have a wider thermal breadth than their hosts (Cohen et al. 2017, Cohen et al. 2020). Our experimental results align with this hypothesis, as we found that ciliate free-living thermal performance curves are left-skewed and parasite infection peaked at lower temperatures than ciliate free-living growth. However, to fully evaluate the hypothesis, data on mosquito thermal performance is needed. Our results also align with a prior observational field study in the *Ae. sierrensis-L. clarki* system, which documented a pattern of increasing parasite prevalence with increasing latitude, consistent with infection peaking at cooler temperatures (Washburn and Anderson 1986).

The ciliate populations varied in the thermal performance of their free-living growth rates, and this variation was consistent with local thermal adaptation. Specifically, we found a marginally significant relationship between peak performance temperature and source temperature (Figure 6). The importance of local adaptation to the environment in shaping host-parasite interactions has long been recognized (Kawecki and Ebert 2004, Sternberg and Thomas 2014). Yet, genotype by environment interactions in parasites are rarely tested. When they are, it is rare that a study includes more than a few populations and connects performance in the experiment to the environmental conditions from which the populations were collected (e.g., Laine 2008, Vale et al. 2008, Mboup et al. 2012, Voyles et al. 2017). To our knowledge, this is the first study to demonstrate a pattern of range-wide local adaptation to temperature in a parasite, in this case in its free-living form; but note that even across 12 populations spanning a 1500 km range, the effect was only marginally significant. This finding demonstrates that the abiotic environment is an important evolutionary force that can strongly influence the fitness of parasites. It also highlights the need to incorporate more than a single population of a parasite in studies of climate and disease.

The temperature-dependent nature of infection varied among parasite populations, indicating that different parasite populations are most infective at distinct temperatures. This result suggests that as we aim to understand the extent of intraspecific variation in parasites, studies conducted at a single temperature may not capture the full extent of variation among populations. Recognizing the complexity of the system, it is important to acknowledge that our experiment overlooked variation in the host, which could contribute to local co-adaptation (Lyberger et al. 2023). Additionally, the environment encompasses more than just annual mean temperature and we were unable to capture factors such as the timing, duration, and intensity of the rainy season, temperature fluctuations, and other habitat features. Conducting this experiment in a field setting with multiple host populations may clarify the role of local (co)adaptation in infection dynamics. It is also of note that in this system, parasitism represents an adaptive response to the threat of predation from the mosquito, and the extent of that threat will vary regionally based on the presence and density of mosquitoes. Coupled with the aforementioned sources of variation, this interplay could drive selection on the ability of parasites to infect hosts across different temperature regimes.

Ultimately, there are relatively few studies on climate change and disease that incorporate intraspecific variation (but see Mitchell et al. 2005, Laine 2008, Vale et al. 2008, Mboup et al. 2012, Gsell et al. 2013, Stevenson et al. 2013, Voyles et al. 2017, Venesky et al. 2022). Most assume that all populations of hosts and parasites respond similarly to temperature. Our experiment provides insight into the extent of variation in temperature-dependent performance between parasite populations in both free-living and infectious traits. Not only was infection temperature dependent, but the relationship between infection and temperature was dependent on parasite population. We also found intraspecific variation in the thermal performance of free-living ciliate growth, and evidence of local adaptation to temperature. Therefore, just as not all areas will experience warming equally, it is unlikely that all populations of hosts and parasites will respond equally to warming. Additionally, our study revealed that infections peaked at a colder temperature than free-living ciliate growth rates, in support of the thermal mismatch hypothesis. This study highlights the importance of considering both host and parasite thermal responses and intraspecific variation when assessing the impacts of climate change on parasites in aquatic ecosystems. We hope this study will motivate further research into the impact of temperature and intraspecific variation on parasite traits, as this will be important for making accurate predictions of disease in a warming climate.

## Acknowledgements

This work was funded by the National Science Foundation (DEB-2011147, with the Fogarty International Center and 2208947 Postdoctoral Research Fellowships in Biology Program), the National Institutes of Health (R35GM133439, R01AI168097, and R01AI102918), the Stanford King Center on Global Development, Woods Institute for the Environment, Center for Innovation in Global Health, the Terman Award, The Rose Hills Foundation, and the Bing-Mooney Fellowship. We would like to thank the staff at Jasper Ridge Biological Preserve, and the many vector control officials who provided invaluable assistance with field collections and lab rearing protocols, including Bret Barner, Nate McConnell, Angie Nakano, Andrew Rivera, and Greg Williams. We would like to thank Allie Lee, Dylan Loth, Isabel Delwel, Mallory Harris, and Desire Nalukwago for their assistance in setting up the experiment, and other members of the Mordecai Lab for their valuable discussions and feedback on the manuscript.

## Declarations

### Author Contributions

KL, SI, and EM originally formulated the idea, KL, JF, and LC conducted fieldwork, KL and SI conducted the experiments and statistical analysis, SI and KL wrote the first draft of the manuscript, and all authors contributed to writing and revising the manuscript.

