## Supplement for "Temperature and intraspecific variation affect host-parasite interactions"

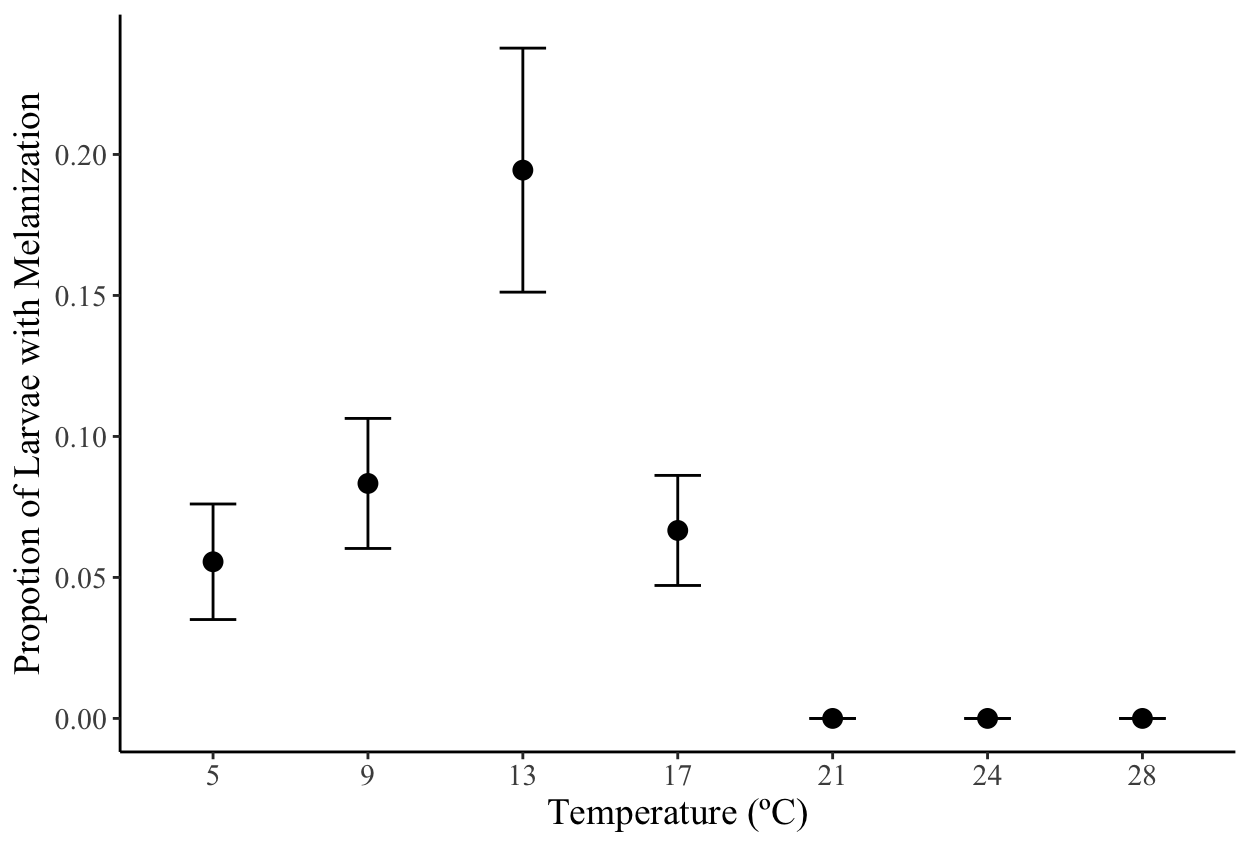

Figure S1. Proportion of larvae with melanization spots per group of 5 larvae (±1 SE) averaged across all parasite populations, shown for each temperature. Melanization peaked at 13°C, and there was no melanization noted at temperatures 21°C, 24°C, and 28°C.

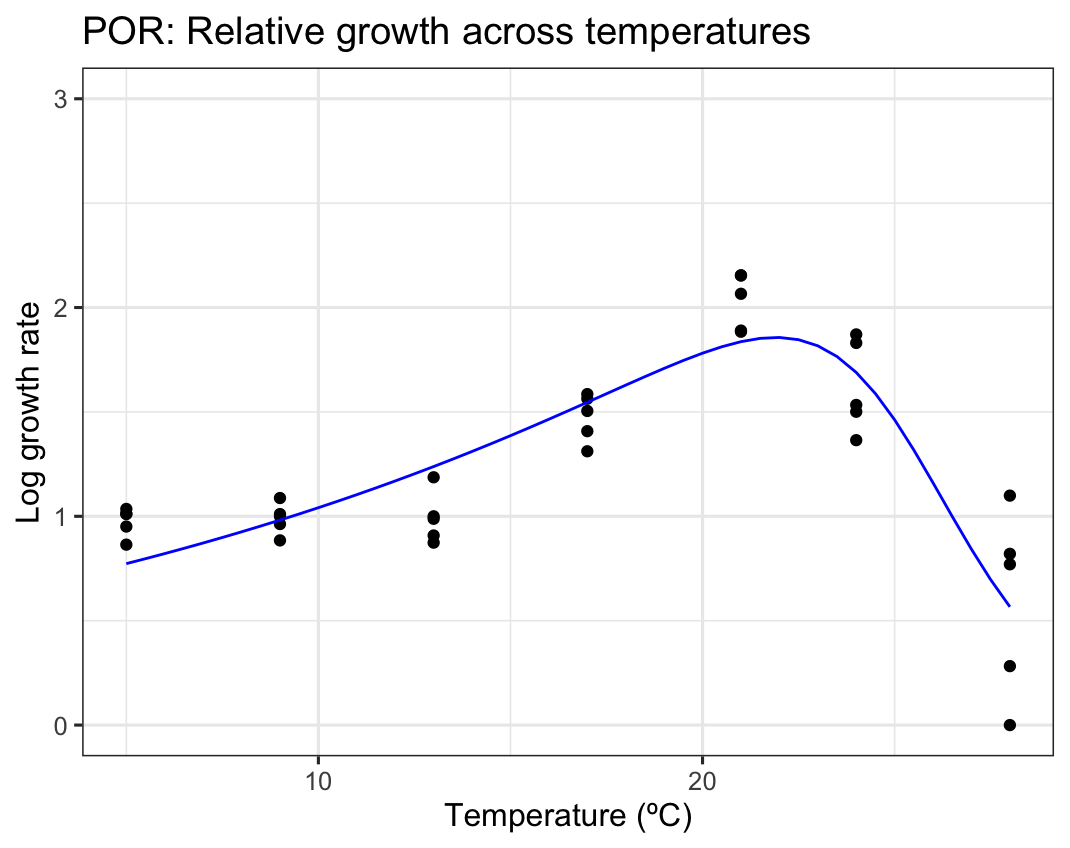

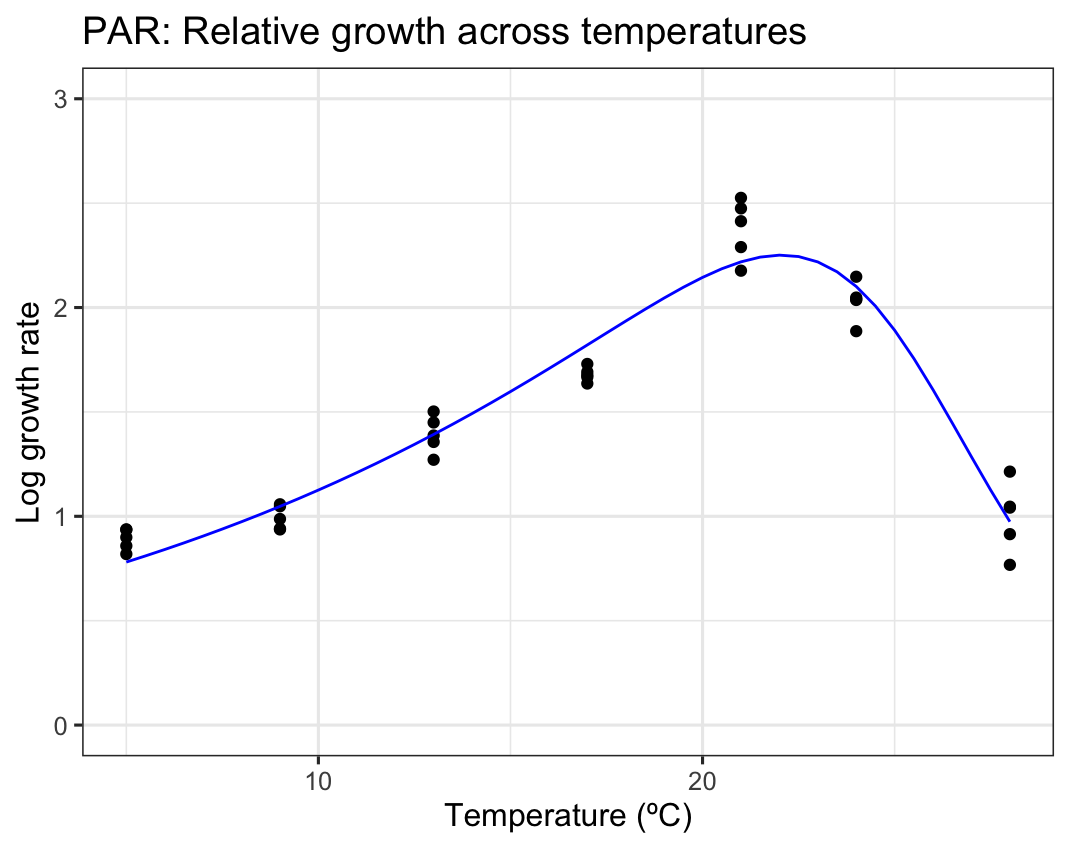

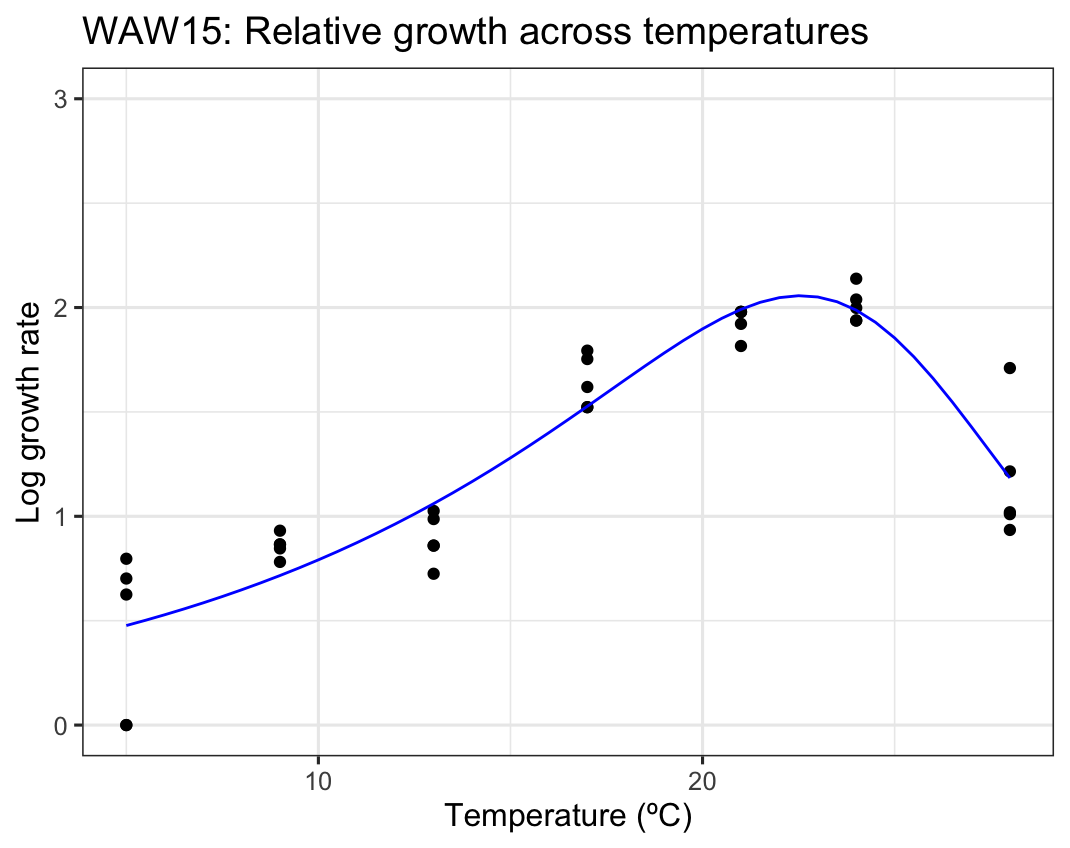

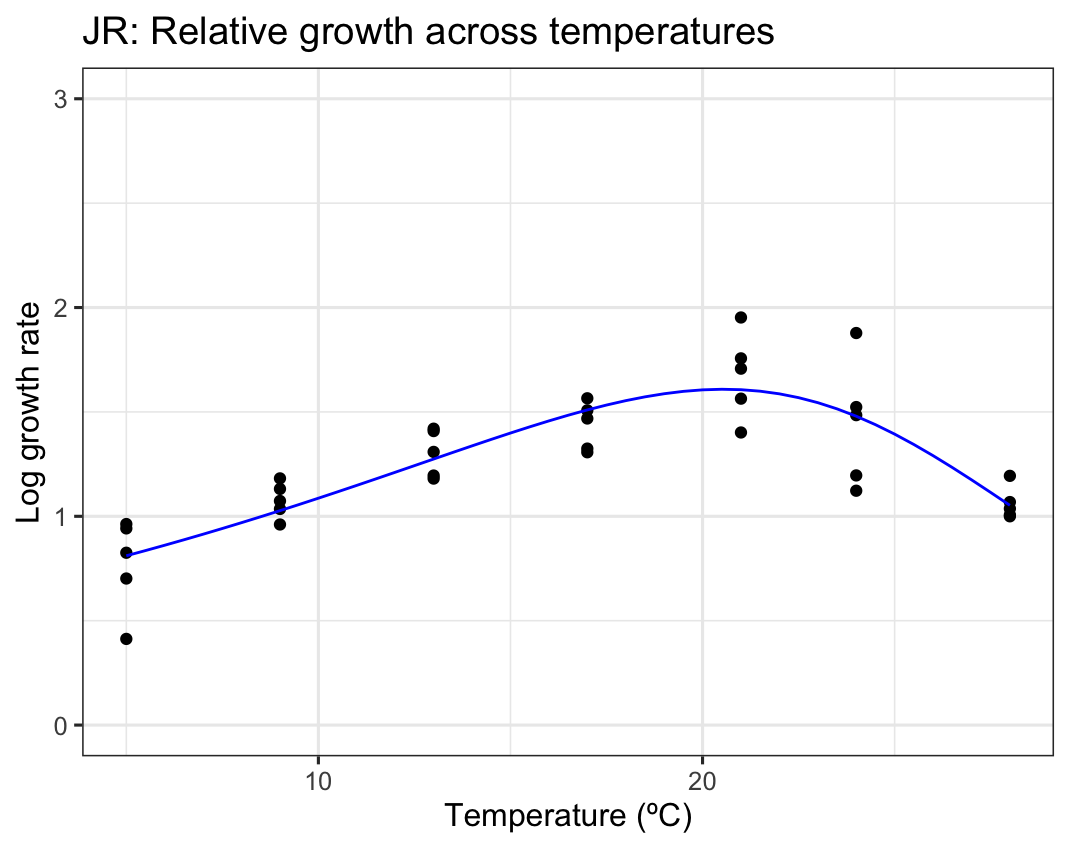

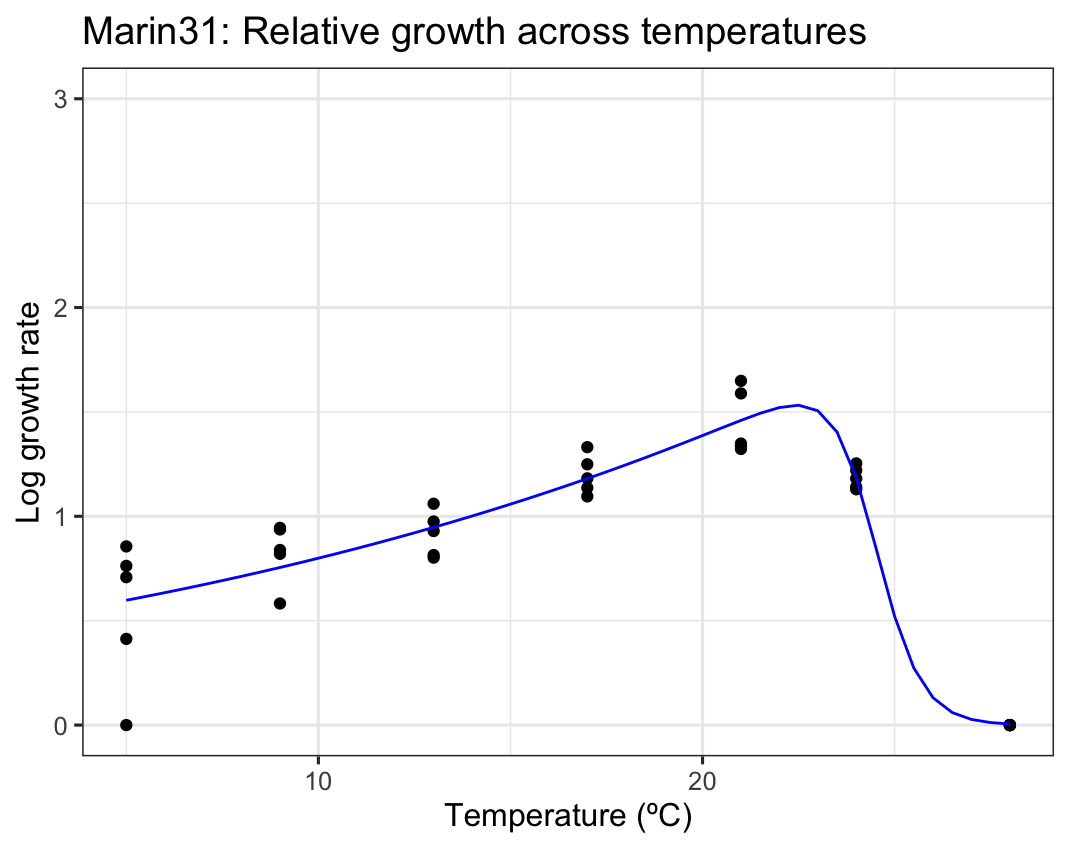

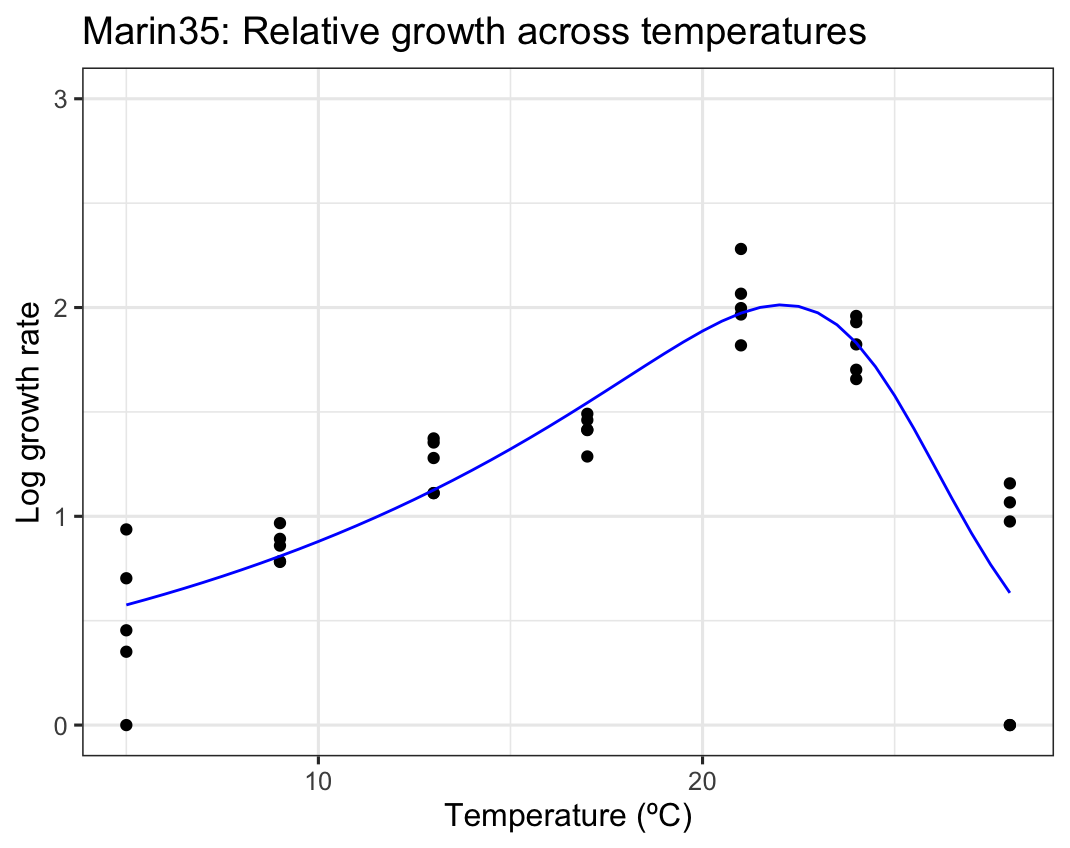

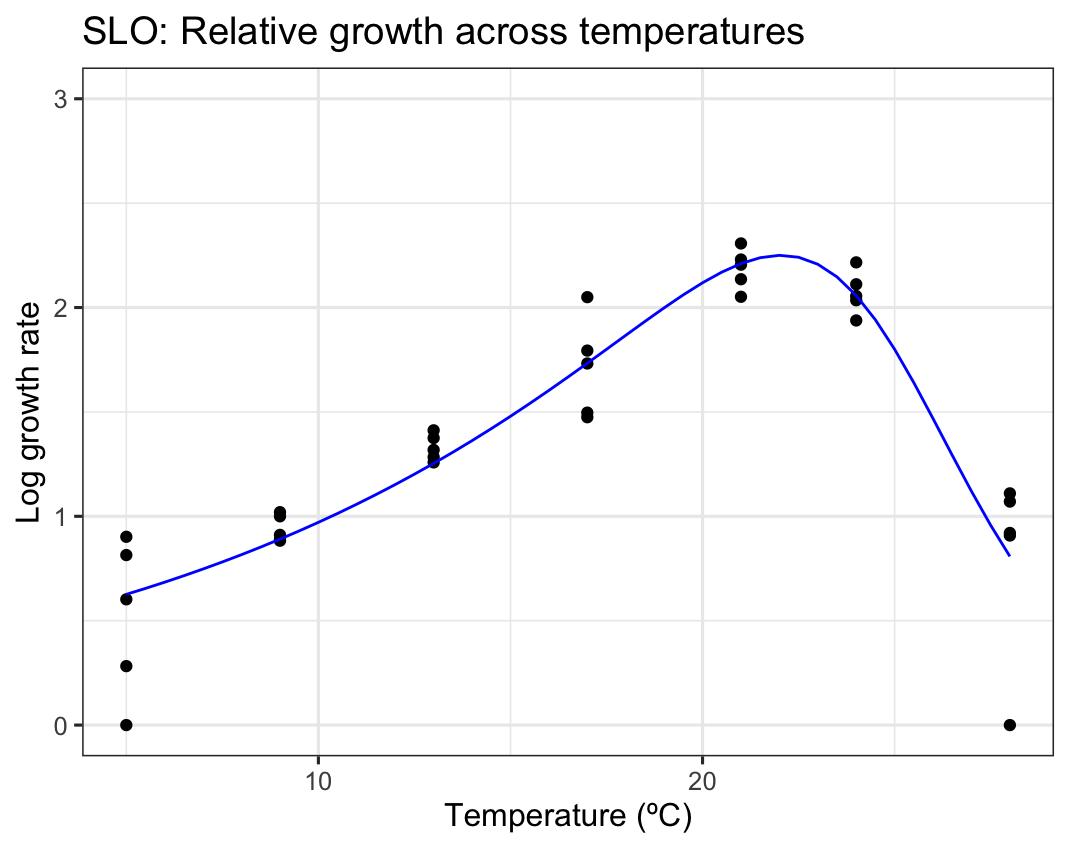

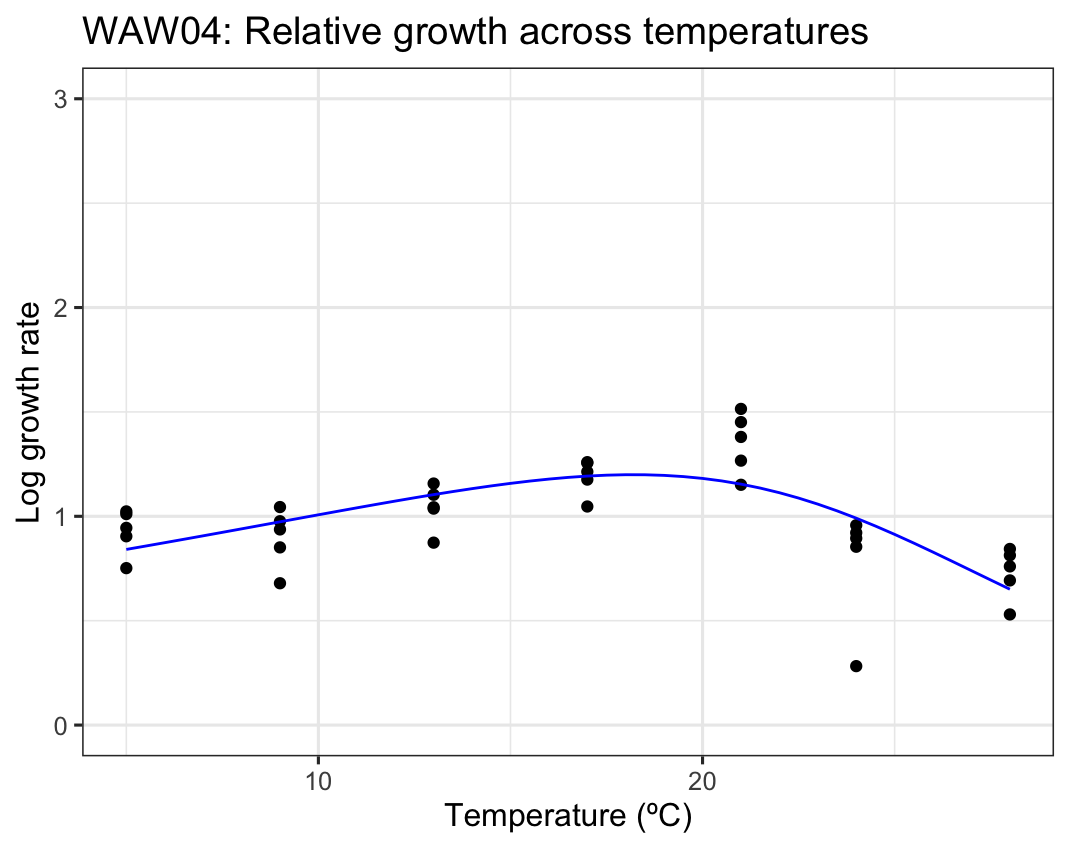

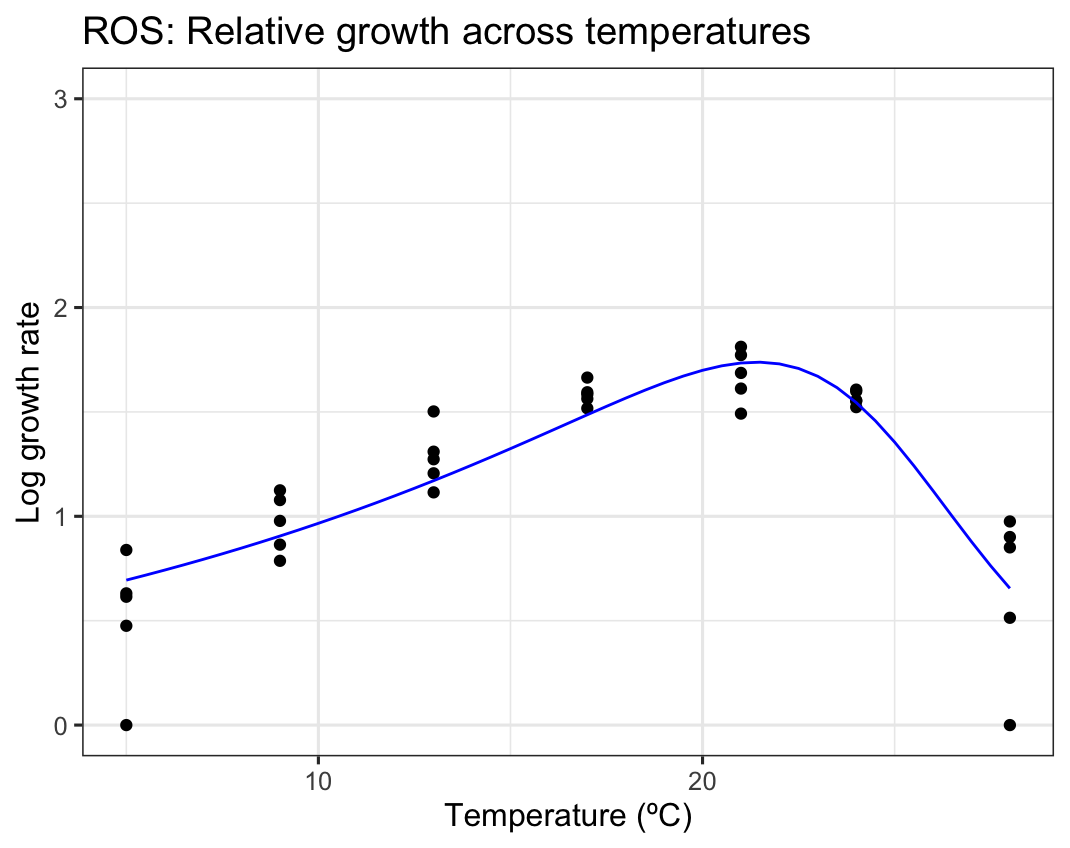

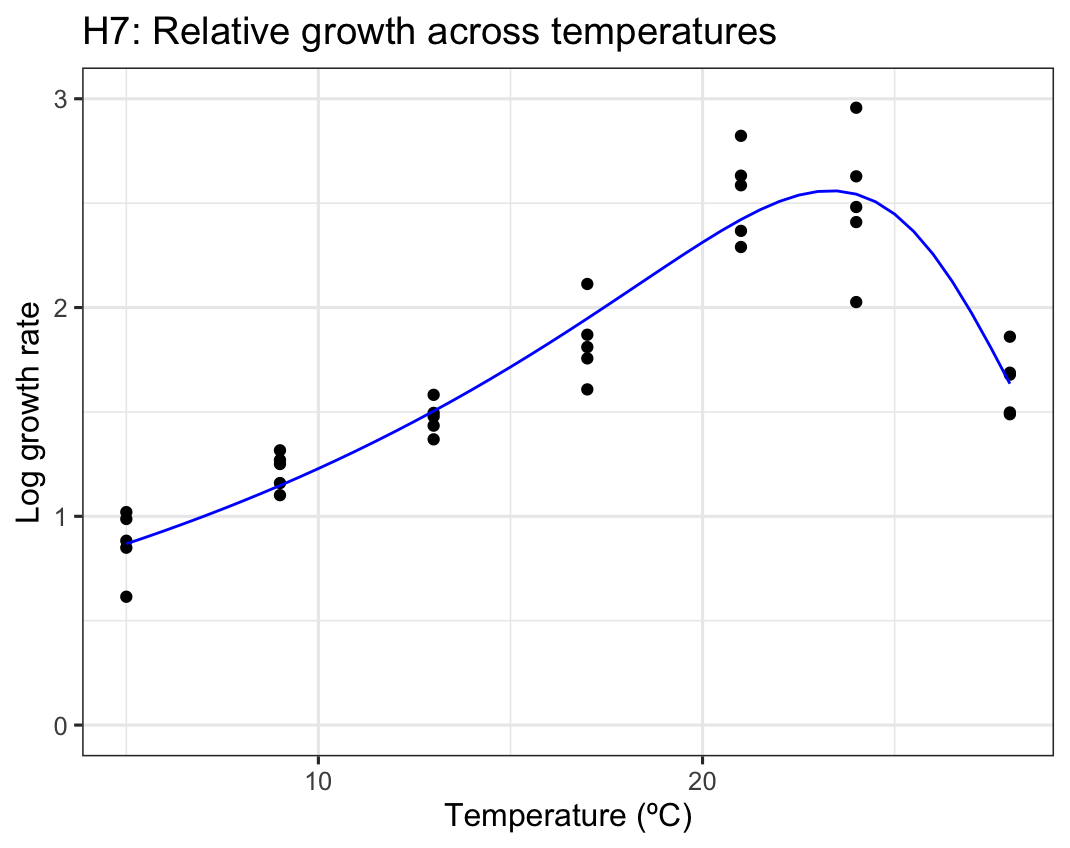

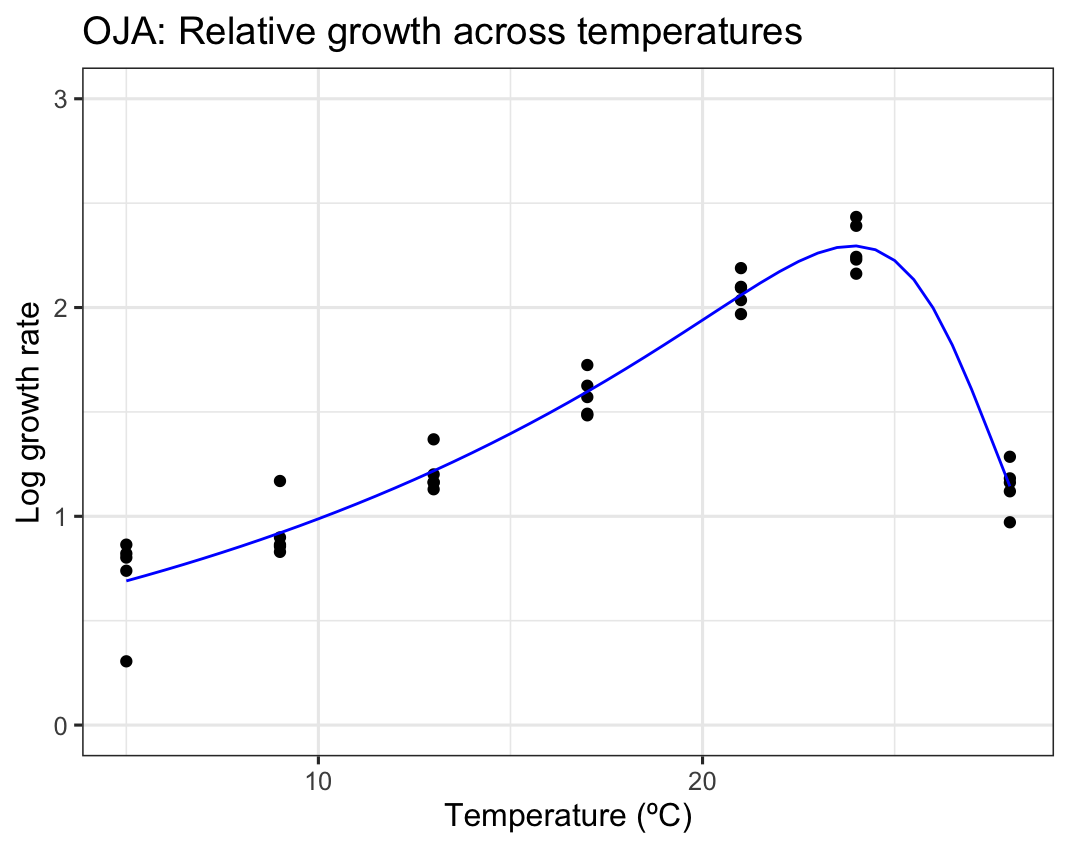

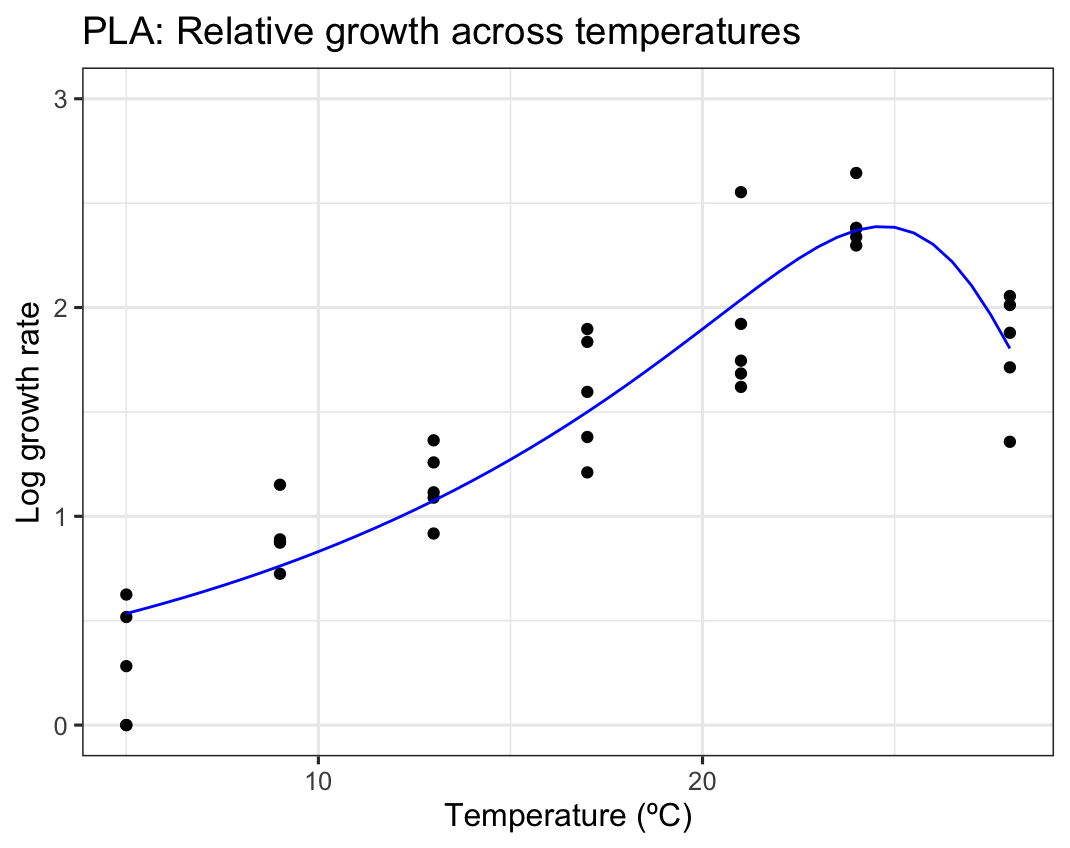
Figure S2. Growth rates of free-living *L. clarki* across temperatures for each population, fit using the Sharpe-Schoolfield model. Growth rates are measured in units of cells per 100ul per day.

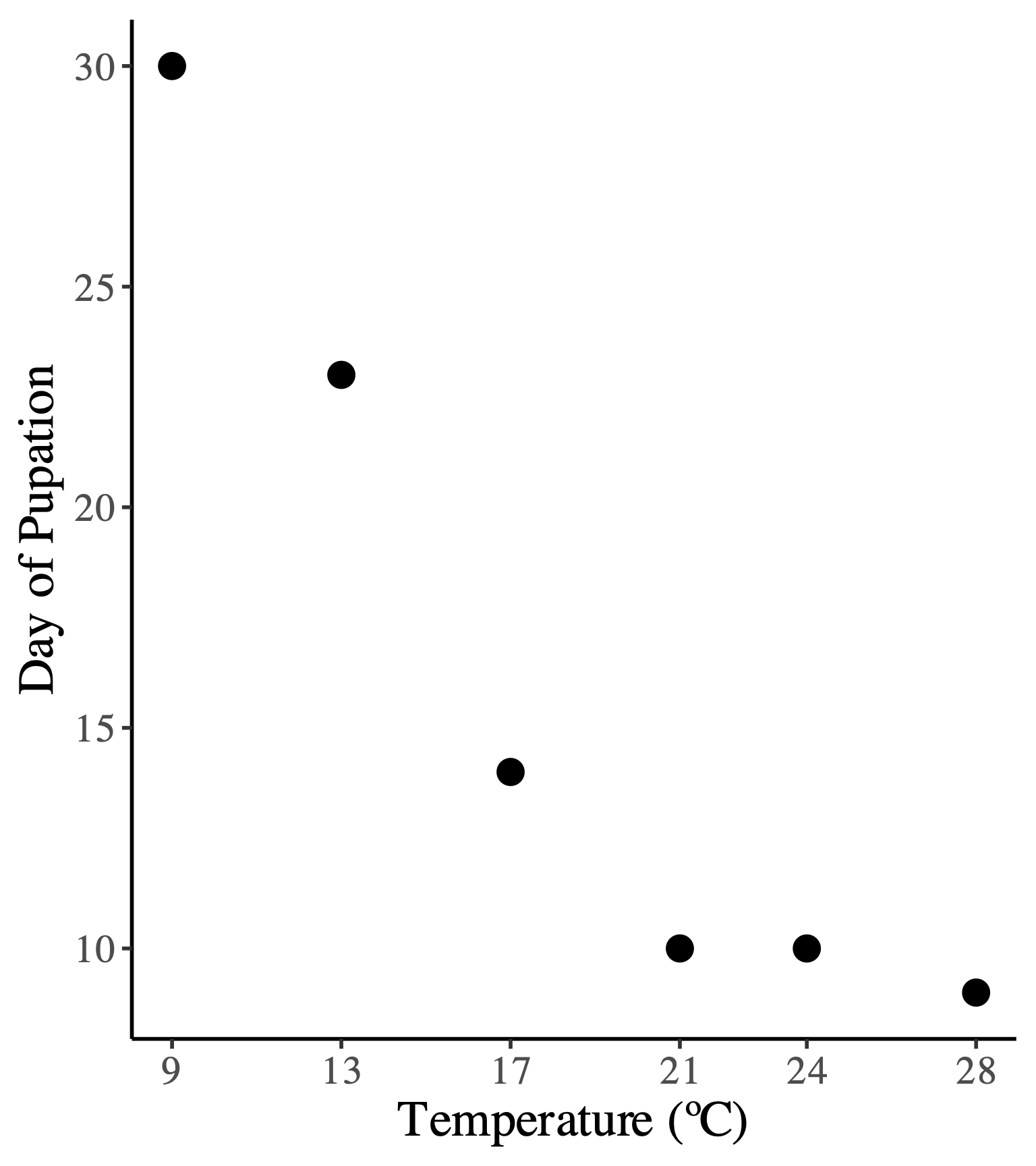

Figure S3. The first day of pupation between the varying populations of *L. clarki* based on temperature. The 5°C population did not pupate.

Table S1. *Lambornella clarki* population localities and average annual temperature.

| Abbreviation | Location | Longitude | Latitude | Source Temperature (°C) | Peak Performance Temperature (°C) |
| --- | --- | --- | --- | --- | --- |
| WAW04 | Wawona, California | 37.5426051 | -119.6271611 | 11.45 | 18.0 |
| ROS | Roseburg, Oregon | 43.2955781 | -122.8923114 | 11.39 | 21.5 |
| POR | Portland, Oregon | 45.446151 | -122.5076021 | 11.26 | 22.0 |
| WAW15 | Oakhurst, California | 37.3925453 | -119.6328531 | 13.44 | 22.5 |
| PLA | Placer, California | 39.0494756 | -120.933641 | 14.47 | 24.5 |
| PAR | Paso Robles, California | 35.65099408 | -120.7353783 | 14.30 | 22.0 |
| Marin31 | Marin County, California | 38.1300867 | -122.5444986 | 15.09 | 22.5 |
| Marin35 | Marin County, California | 38.1304913 | -122.543991 | 15.09 | 22.0 |
| JR | Jasper Ridge, California | 37.40912 | -122.23321 | 14.06 | 20.5 |
| SLO | San Luis Obispo County, California | 35.2642605 | -120.7178764 | 14.29 | 22.0 |
| OJA | Ojai, California | 34.3567726 | -119.3155263 | 15.85 | 24.0 |
| H7 | Hollenbeck Canyon, California | 32.68192291 | -116.8186111 | 17.82 | 24.0 |

Table S2. Correlations between peak thermal performance temperature of free-living *L. clarki* and 19 bioclimatic variables at natal source location.

| Climate Variable | Correlation (r-value) |
| --- | --- |
| Annual mean temperature | 0.65 |
| Mean Diurnal Range (Mean of monthly (max temp - min temp)) | 0.14 |
| Isothermality (BIO2/BIO7) (×100) | 0.19 |
| Temperature Seasonality (standard deviation ×100) | -0.17 |
| Max Temperature of Warmest Month | 0.31 |
| Min Temperature of Coldest Month | 0.51 |
| Temperature Annual Range (BIO5-BIO6) | -0.06 |
| Mean Temperature of Wettest Quarter | 0.52 |
| Mean Temperature of Driest Quarter | 0.53 |
| Mean Temperature of Warmest Quarter | 0.53 |
| Mean Temperature of Coldest Quarter | 0.52 |
| Annual Precipitation | -0.20 |
| Precipitation of Wettest Month | -0.21 |
| Precipitation of Driest Month | -0.14 |
| Precipitation Seasonality (Coefficient of Variation) | 0.26 |
| Precipitation of Wettest Quarter | -0.19 |
| Precipitation of Driest Quarter | -0.15 |
| Precipitation of Warmest Quarter | -0.14 |
| Precipitation of Coldest Quarter | -0.20 |
